# Hierarchical Interplay between H3K27ac and H3K4me3 in Transcriptional Regulation

**DOI:** 10.64898/2026.05.11.724317

**Authors:** Chenwei Zhou, Chanjuan Dong, Weiye Zhao, Fu-Sen Liang

## Abstract

H3K27ac and H3K4me3 are enriched at transcriptional start sites and have been implicated in transcription. However, how these marks concertedly regulate transcription is not fully understood. Here, we developed a dual chemically inducible CRISPR/dCas9-based epigenome editing system that enables independent, temporal and transcription stage-specific modulation of H3K27ac and H3K4me3 at a specific gene locus. Stage-specific removal of H3K4me3 impaired RNA polymerase II recruitment, increased promoter-proximal pausing, reduced productive elongation, and accelerates mRNA decay via increased m^6^A deposition. Losing both H3K27ac and H3K4me3 rapidly abolished transcriptional activity, while preserving H3K4me3 without H3K27ac can partially sustain transcription. These findings revealed a functional hierarchy and interdependence between H3K27ac and H3K4me3 in different transcription stages and the established versatile tool will contribute to the functional dissection of the temporal dynamics of chromatin modifications in gene regulation.

## MAIN

Regulation of gene expression is fundamental to cellular function and their responses to environmental cues.^1,2^ A key underlying mechanism is histone modification-mediated remodeling of chromatin that shape DNA accessibility and transcriptional activity.^3–5^ Among these modifications, histone H3 lysine 27 acetylation (H3K27ac) and histone H3 lysine 4 trimethylation (H3K4me3) are highly enriched at active promoters and frequently colocalize, suggesting a functional synergy in transcription.^6–10^ H3K27ac is known to remodel chromatin structure and facilitate the recruitment of transcriptional machinery, including RNA polymerase II, thereby promoting transcription initiation.^11–14^ Moreover, H3K27ac-associated bromodomain proteins such as bromodomain-containing protein 4 (BRD4) recruit positive transcription elongation factor b (p-TEFb), which phosphorylates RNA polymerase II (pol-II) and promotes its pause-release.^15–17^ H3K4me3 has been recognized as a hallmark of transcription initiation through the recruitment of plant homeodomain (PHD)-containing proteins such as TBP-associated factor 3 (TAF3).^18–20^ More recently, H3K4me3 has been shown to recruit integrator complex subunit 11 (INTS11) that evicts paused RNA pol-II and promotes productive elongation.^19,21^ Ectopic H3K4me3 writing studies have yielded conflicting results regarding whether H3K4me3 is sufficient to induce transcription.^22,23^

CRISPR-based epigenome editing technologies have enabled programmable, locus-specific manipulation of chromatin states. Nuclease-dead Cas9 (dCas9) has been fused to chromatin-modifying effectors to achieve sequence-specific editing of histone modifications.^24–27^ To interrogate the functional and temporal interplay between H3K27ac and H3K4me3 during transcription, we developed a dual chemically inducible CRISPR/dCas9-based epigenome editing system that integrates orthogonal chemically-induced proximity (CIP) modules within a dCas9 targeting framework,^28–31^ which enables rapid, reversible, and stage-specific independent modulation of H3K27ac and H3K4me3 at defined genomic loci with a minute-scale temporal resolution. Using this system, we showed that the distinct and cooperative roles of H3K27ac and H3K4me3 in transcriptional activation. H3K27ac is indispensable for transcription initiation and maintenance and H3K4me3 sustains maximal transcriptional output by promoting efficient elongation and enhancing mRNA stability.

## RESULTS

### H3K4me3 Is Required for Optimal Transcription Initiated by H3K27ac

To determine the contribution of H3K4me3 to H3K27ac-driven gene expression, we used a reported dCas9-P300(CD) acetyltransferase fusion protein to deposit H3K27ac at the IL1RN promoter (Extended Data Fig. 1a).^32^ We cloned a fusion protein of dCas9 and the catalytic core domain of H3K4me3/me2 demethylase KDM5B to remove H3K4me3 (Extended Data Fig. 1b).^41–44^ We transfected HEK293T cells with either dCas9-P300(CD), dCas9-PYL (negative control^23,37^), or both dCas9-P300(CD) and dCas9-KDM5B(CD), with four sgRNAs targeting the IL1RN promoter region (Extended Data Fig. 1c).^32^ Chromatin immunoprecipitation (ChIP) and quantitative polymerase chain reaction (qPCR) assays were used to quantify the enrichment of H3K27ac and H3K4me3 near transcription start site (TSS). We observed that dCas9-P300(CD) alone increased both H3K27ac and H3K4me3 levels at the targeted IL1RN promoter,^23^ while co-expression of dCas9-P300(CD) and dCas9-KDM5B(CD) resulted in complete depletion of H3K4me3 without affecting H3K27ac (Extended Data Fig. 1d). We quantified IL1RN mRNA levels by reverse transcription qPCR (RT-qPCR) and observed that while H3K27ac alone led to robust activation of IL1RN transcription, the simultaneous removal of H3K4me3 caused a significant reduction in the IL1RN mRNA level but not completely abolished (Extended Data Fig. 1e). These results indicate that H3K4me3 plays a key role in supporting transcription.

### Establishing a Dual Inducible Editing System for Independent H3K27ac and H3K4me3 Editing

To investigate the regulatory relationship of H3K4me3 and H3K27ac during the time course of transcription, we developed a dual chemically inducible CRISPR-based epigenome editing system integrating orthogonal abscisic acid (ABA)-inducible ABI-PYL and rapamycin (Rap)-inducible FRB-FKBP CIP systems to independently and temporally modulate H3K27ac and H3K4me3 (Fig. 1a). H3K27ac writing was achieved by fusing ABI with P300(CD) and a Flag tag, while H3K4me3 removal was mediated by fusing FRB with KDM5B(CD) and a HA tag.^29,30^ Genome site-specific targeting was accomplished by fusing dCas9 with FKBP, PYL and a V5 tag. We cloned 10 varying fusion configurations of dCas9, PYL and 3×FKBP driven either by a weak PGK promoter (V1-ID1-5) or a strong CAG promoter (V2-ID1-5) (Extended Data Fig. 2a).^38^ We also cloned the recruitable effectors Flag-ABI-P300(CD) (V1-ID6), and HA-FRB-KDM5B(CD) (V1-ID7) driven by the PGK promoter (Extended Data Fig. 2a). Using IL1RN mRNA expression as a screening criterion, we found that several configurations (V1-ID2, V1-ID4, V2-ID2, V2-ID4, V2-ID5) enabled robust gene activation upon ABA addition (H3K27ac writing) but did not reduce expression levels when both ABA and Rap were added (with H3K27ac but removing H3K4me3) (Extended Data Fig. 2b), suggesting ineffective H3K4me3 erasure and/or HA-FRB-KDM5B(CD) recruitment.

**Fig. 1.**
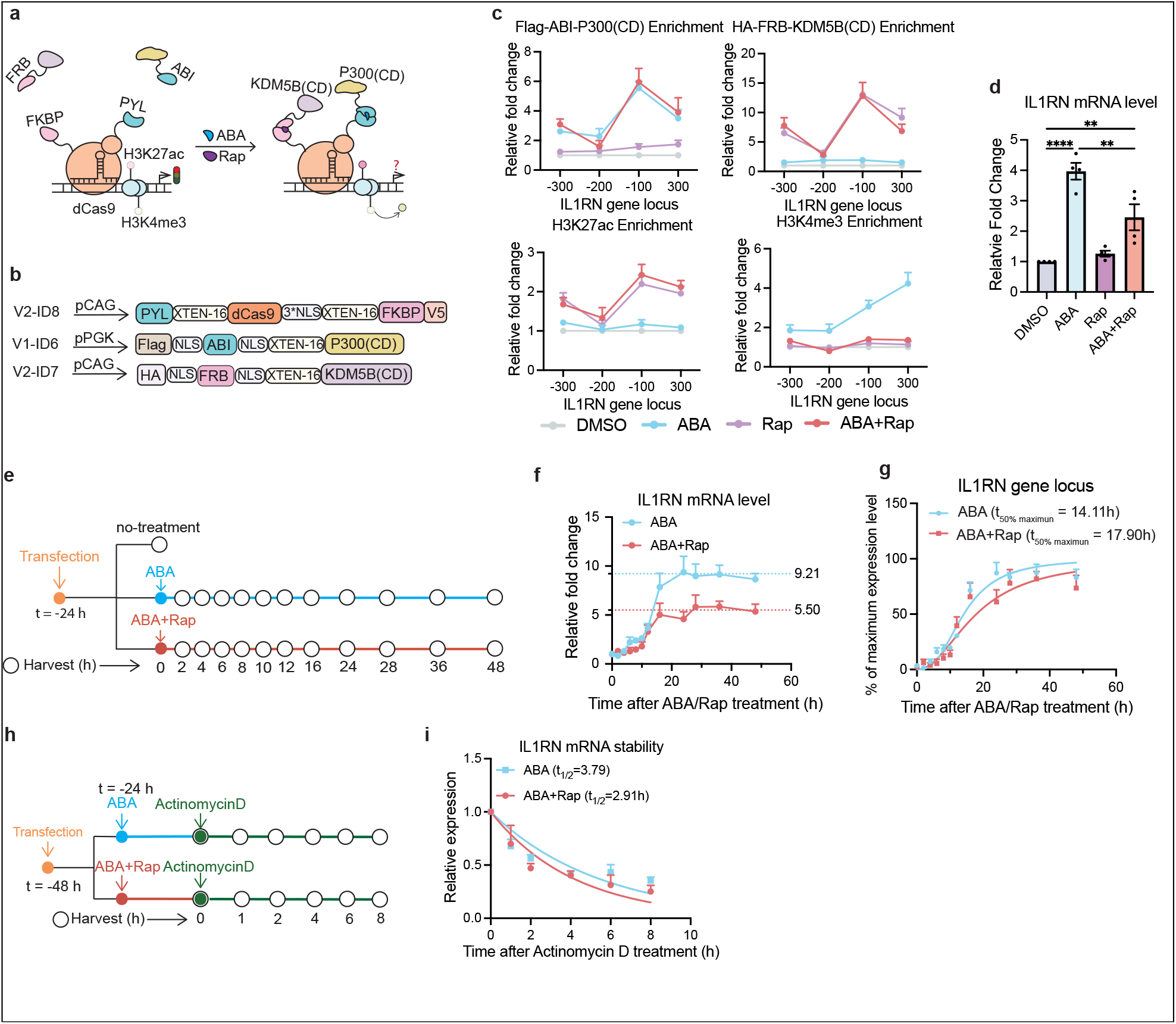
Establishment of a dual-inducible epigenome editing system reveals H3K4me3’s role in accelerating transcription and stabilizing mRNA during H3K27ac-induced activation at the IL1RN gene locus. **a**, The dual-inducible H3K27ac and H3K4me3 editing system enabling independent recruitment of P300(CD) and KDM5B(CD) using ABA- and Rap-inducible dimerization modules. **b**, Constructs of optimized dual editing system used in this study. **c**, ChIP-qPCR analysis of Flag-ABI-P300(CD), HA-FRB-KDM5B(CD), H3K27ac, and H3K4me3 enrichment at the IL1RN promoter following treatment with DMSO, ABA, Rap, or ABA+Rap. **d**, IL1RN mRNA levels determined by RT-qPCR under the indicated conditions. **e**, Time-course experiments to determine IL1RN mRNA levels from collected cells treating with ABA or ABA+Rap at indicated time points. **f**, IL1RN mRNA accumulation measured by RT-qPCR after ABA (steady-state mRNA level: 9.21) or ABA+Rap treatment (steady-states mRNA level: 5.50). **g**, Nonlinear regression analyses of IL1RN expression rate. t_50% maximum_ is defined as the time reaches 50% maximum expression level (R^2^: 0.90 for ABA treatment; 0.82 for ABA+Rap treatment). **h**, Time-course experiments to determine mRNA stability by quantifying IL1RN mRNA levels using RT-qPCR at indicated time points after actinomycin D treatment. **i**, The decay curves of IL1RN mRNA after actinomycin D treatment. t_1/2_ is defined as the mRNA half-life (R^2^: 0.67 for ABA treatment; 0.66 for ABA+Rap treatment). The enrichment fold changes were calculated by comparing to the DMSO-treated samples in (c) and (d), to o samples at t = 0 h (without treatment) in (f), and to the expression level under ABA or ABA+Rap condition before Actinomycin D treatment (t= 0 h) in (h). Data represent mean ± s.e.m. (n= 4). Statistical significance was determined by two-way ANOVA (c) or one-way ANOVA (d). ns, P > 0.5; *P ≤ 0.05; **P ≤ 0.01; ***P ≤ 0.001; ****P ≤ 0.0001. The detailed statistical analysis data (p-values) can be found in Supplementary Table **1**.

We suspect that the abundant endogenous FKBP proteins may compete with the FKBP-fusion proteins for HA-FRB-KDM5B(CD) binding.^39,40^ As a result, we clone HA-FRB-KDM5B(CD) driven by stronger promoters, including pCAG (V2-ID7) and pCMV (V3-ID7) (Extended Data Fig. 3a).^38^ Combining V2-ID4 (dCas9-PYL-3×FKBP) with V1-ID6 (Flag-ABI-P300(CD)), and V1/2/3-ID7 (HA-FRB-KDM5B(CD) with different promoters), we observed that HA-FRB-KDM5B(CD) driven by the high-expression promoters (V2-ID7 or V3-ID7) significantly reduced gene expression upon ABA+Rap treatment (Extended Data Fig. 3b), indicating successful H3K4me3 removal. ChIP-qPCR analyses confirmed that adding ABA alone effectively recruited Flag-ABI-P300(CD) and increased H3K27ac and H3K4me3 enrichment (Extended Data Fig. 3c). Rap alone resulted in efficient recruitment of HA-FRB-KDM5B(CD) with no effects on H3K27ac or H3K4me3 (Extended Data Fig. 3c). ABA+Rap treatment led to significant enrichment of Flag-ABI-P300(CD), HA-FRB-KDM5B(CD), and H3K27ac, with near complete depletion of H3K4me3 (Extended Data Fig. 3c). However, both Flag-ABI-P300(CD) and H3K27ac levels were significantly reduced compared to ABA alone, with no significant decrease in HA–FRB– KDM5B(CD) recruitment (Extended Data Fig. 3c), indicating a disruption of Flag-ABI-P300(CD) recruitment and H3K27ac editing, potentially due to the spatial interference from the co-recruited HA-FRB-KDM5B(CD).

To address this issue, we decreased the number of FKBP from 3 copies to one (V2-ID8, Fig. 1b) to reduce the number of HA-FRB-KDM5B(CD) being recruited. We then tested the H3K27ac and H3K4me3 co-editing using V2-ID8 combined with V1-ID6 and V2-ID7 and found that when both ABA and Rap were applied, both Flag-ABI-P300(CD) and HA-FRB-KDM5B(CD) were efficiently recruited, with a robust increase level of H3K27ac while and a complete depletion of H3K4me3 (Fig 1c), which also reduced IL1RN expression (Fig. 1d). These results indicated that we successfully optimized the H3K27ac/H3K4me3 dual editing system for the following studies.

### Loss of H3K4me3 Reduces the Rate of Transcription and the Stability of mRNA

The dual-inducible epigenome editing system developed above was used to monitor H3K27ac-induced gene expression time course in the presence or absence of H3K4me3. HEK293T cells co-transfected with dual-inducible modules (V1-ID6, V2-ID8 and V2-ID7) and IL1RN sgRNAs for 24 h were then treated with ABA, ABA+Rap, or DMSO, and harvested at the indicated time points within 48 h post-treatment (Fig. 1f), followed by RT-qPCR analyses. Compared to cells treated with ABA alone (gained both H3K27ac and H3K4me3), we observed that cells treated with both ABA and Rap (gained H3K27ac but depleting H3K4me3) had a reduced steady-state mRNA level by 40% compared to ABA treated cells (Fig. 1g). Similar experiments were performed at GRM2 and MYOD1 gene loci (Extended Data Fig. 4a and 4e) and the comparable reduction of steady-state mRNA levels from depleting H3K4me3 were also observed (30% for MYOD1 and 35% for GRM2) (Extended Data Fig. 4b and 4f).

A reduced steady-state mRNA level can potentially be resulted from a decreased RNA synthesis rate.^41^ To determine whether H3K4me3 influences transcription speed, we analyzed transcription kinetics and calculated the time required to reach 50% of maximum expression (t_50% maximum_) using nonlinear regression analyses. At the IL1RN locus, ABA-treated cells reached 50% of maximum expression at 14.11 h, whereas cells treated with both ABA and Rap took 17.90 h, a 27% reduction in mRNA production rate (Fig. 1g). Comparable delays were observed for MYOD1 (t_50% maximum_ = 9.72 h for ABA treatment and 11.92 h for ABA+Rap treatment, a 23% reduction) and GRM2 (t_50% maximum_ = 10.44 h for ABA treatment and 12.45 h for ABA+Rap treatment, a 19% reduction) (Extended Data Fig. 4c and 4g).

Steady-state mRNA abundance reflects the balance between mRNA synthesis and degradation.^42^ To assess if H3K4me3 also influences mRNA degradation, HEK293T cells were co-transfected with dual-inducible modules and either IL1RN, GRM2 or MYOD1 sgRNAs for 24 h, followed by treating with ABA alone, ABA+Rap, or DMSO for another 24 h. Actinomycin D, an inhibitor for transcription,^42^ was then added to block new RNA synthesis, and cells were harvested at multiple time points after treatment (Fig. 1h) and subjected to RT-qPCR assays to measure the mRNA half-life under each condition. We observed that the half-life of IL1RN mRNA was 3.79 h in ABA-treated cells and 2.91 h in cells treated with ABA+Rap (Fig. 1i), indicating a 23% faster decay rate when H3K4me3 was depleted. Similar increases in decay rate were observed for MYOD1 (5.38 h for ABA treatment and 3.97 h for ABA+Rap treatment, a 26% faster decay rate) and GRM2 (11.03 h for ABA treatment; 7.49 h for ABA+Rap treatment, a 32% faster decay rate) (Extended Data Fig. 4d and 4h).

Overall, these results showed that H3K4me3 is not only required to maintain the full rate of mRNA synthesis but also regulates the stabilization of transcribed mRNAs, highlighting its dual role in supporting robust gene expression both transcriptionally and post-transcriptionally.

### H3K27ac Triggers Transcription Initiation and Sequential Recruitment of Transcriptional Machineries

Transcription proceeds rapidly through distinct kinetic stages. Initiation occurs within minutes of promoter activation by recruiting RNA pol-II and its phosphorylation at the Ser5 of C-terminal domain (CTD) (RNA pol-II Ser5Ph),^43–45^ followed by promoter-proximal pausing within minutes stabilized by Negative Elongation Factor (NELF),^46–48^ where RNA pol-II remains stalled for several minutes before being released into productive elongation by positive transcription elongation factor b (p-TEFb), which phosphorylates NELF, and RNA pol-II CTD at Ser2 (RNA pol-II Ser2Ph).^49,50^ Once past nucleosomal barriers, elongation completes within 5-30 minutes depending on gene length.^44,51^ H3K4me3 has been shown to recruit transcription regulators to facilitate transcription initiation and RNA pol II promoter proximal pause-release.^18,21^

To investigate if H3K4me3 affect any of these stages, we temporally depleted H3K4me3 at specific transcription stages after gene activation induced by H3K27ac. The CIP-integrated epigenome editing methods have been shown to be fast and able to remodel chromatin within minutes.^52,53^ To confirm the fast kinetics of our editing system, we characterized the kinetics of ABA-induced recruitment of Flag-ABI-P300(CD), the installation of H3K27ac, and the resulting increase of H3K4me3. HEK293T cells were co-transfected with Flag-ABI-P300(CD), PYL-dCas9-FKBP-V5, and IL1RN-targeting sgRNAs for 48 h, then treated with ABA and harvested cells over a short time course (5, 15, 30 and 60 min after ABA addition, Fig. 3a and Extended Data Fig. 5a). ChIP-qPCR analyses revealed the rapid recruitment of Flag-ABI-P300(CD) to the promoter within 5 min (Extended Data Fig. 5b), coinciding with immediate H3K27ac deposition within 5 min at the locus (Extended Data Fig. 5b). Notably, H3K4me3 enriched within 15 min post-ABA addition (Extended Data Fig. 5b), indicating that H3K4me3 writing occurred immediately following initial H3K27ac deposition.

We next mapped the timing and order of deposition and recruitment of H3K27ac, H3K4me3 and stage-specific transcriptional regulators along the gene activation time course. According to established models, initiation and promoter-proximal pause factors such as RNA pol-II Ser5P and NELF localized near the TSS and shifted downstream as elongation-associated factors including p-TEFb and RNA pol-II Ser2Ph progressively accumulated, reflecting the transition from initiation to productive elongation.^54–56^ To capture this spatial progression, we selected primer sites 100 bp upstream of the TSS and 100, 200, and 300 bp downstream to monitor initiation, pausing, pause-release, and elongation, respectively. We transfected cells with Flag-ABI-P300(CD), PYL-dCas9-FKBP-V5, and IL1RN-targeting sgRNAs for 48 h, then treated cells with ABA over the indicated time course before analyzed by ChIP-qPCR for the enrichment of H3K27ac, H3K4me3 and stage-specific regulator around TSS along the time course (Fig. 2a and 2b). We defined a significant increase at each time point and at each locus when the p-value (calculated from the two-way ANOVA statistical analyses) for the enrichment fold change under each condition was < 0.05 when compared to no ABA treatment. We found that H3K27ac was significantly enriched within 5 min following ABA treatment at both upstream (−100 bp, p = 0.0195) and downstream (+100 bp: p = 0.0372; +200 bp: p = 0.0369; +300 bp: p = 0.035) of the TSS (Fig. 2c and Supplementary Table 1). H3K4me3 deposition reached significance at 10 min at the −100 bp position (p = 0.0445) but not at other sites at this time point (+100 bp: p = 0.998; +200 bp: p = 0.1954; +300 bp: p = 0.0534). By 15 min, all positions displayed significant enrichment (all p ≤ 0.0218) (Fig. 2c and Supplementary Table 1). To define the onset of each transcription stage, we identified it based on the earliest time point showing significant enrichment of the corresponding stage-specific transcription regulators. The initiation marker RNA pol-II Ser5Ph was significantly enriched by 10 min at −100 bp (p = 0.0162) and +100 bp (p = 0.0054), while the enrichment at downstream sites was not detected at this time (+200 bp: p = 0.6530; +300 bp: p = 0.7788) (Fig. 2c and Supplementary Table 1). Promoter-proximal pause marker NELF was enriched at +100 bp by 15 min (p = 0.0227) but not at other sites at 15 min (−100 bp: p = 0.3637; +200 bp: p = 0.1563; +300 bp: p = 0.2916) (Fig. 2c and Supplementary Table 1). Pause-release and productive elongation marker p-TEFb was first enriched at +100 bp at 20 min (p = 0.0136) and subsequently at +200 bp (p = 0.0011) and +300 bp (p = 0.0182) by 30 min (Fig. 2c and Supplementary Table 1). Consistent with this, the elongation marker RNA pol-II Ser2Ph was detected at +100 bp at 20 min (p = 0.036) and shifted downstream to +200 bp (p = 0.0222) and +300 bp (p = 0.0285) by 30 min (Fig. 2c and Supplementary Table 1). The temporal lag in p-TEFb and RNA pol II Ser2Ph enrichment between +100 bp and +200 bp is consistent with the time required for pol II to traverse the initial nucleosome barrier.^54–56^ We further applied nonlinear regression analyses using ChIP-qPCR time-course data to estimate the time required to reach 50% maximal enrichment (t_50% maximum_) for each modification and regulator. H3K27ac was the first to accumulate (t_50% maximum_= 9.22 min at -100 bp), followed by RNA pol-II Ser5Ph (t_50% maximum_= 12.45 min at -100 bp) (Fig. 2d). H3K4me3 followed initiation (t_50% maximum_= 16.04 min at -100 bp and 19.54 min at +100 bp) and was closely coincident with NELF recruitment (t_50% maximum_= 18.81 min at +100 bp) (Fig. 3d). Pause-release and elongation followed with p-TEFb and RNA pol-II Ser2Ph reaching 50% enrichment at +100 bp around 28 min and delayed accumulation downstream (t_50% maximum_ = 41.14 - 47.73 min at +200/+300 bp) (Fig. 2d). Overall, the observed timeline of H3K27ac and H3K4me3 deposition as well as the recruitment sequence of transcription stage markers closely mirror the canonical transcription cycle, providing a precise temporal map of ABA-induced transcriptional events at the IL1RN locus.

**Fig. 2.**
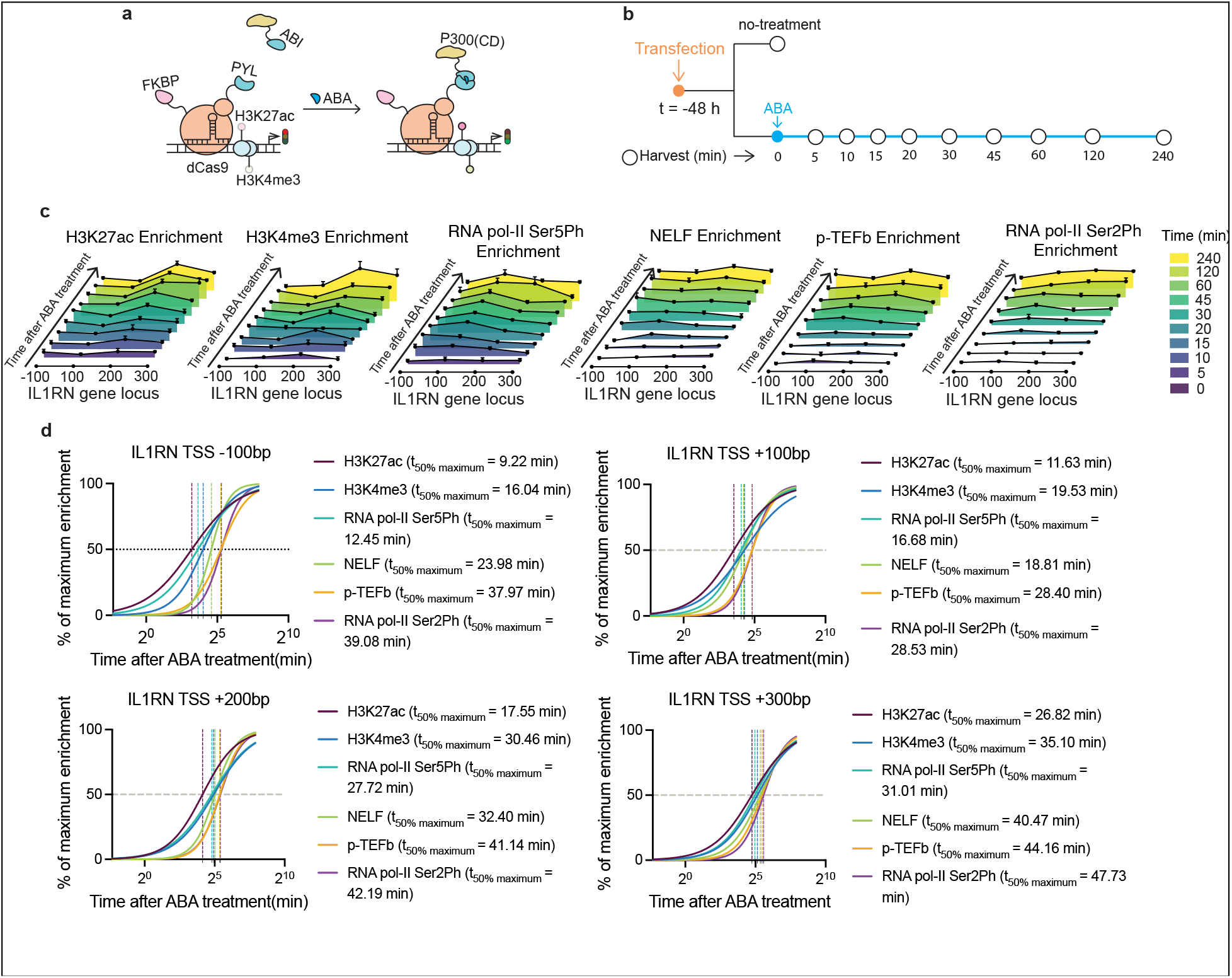
H3K27ac deposition primes transcriptional initiation and sequential recruitment of transcriptional regulators at the IL1RN Locus. **a**, ABA-induced recruitment of Flag-ABI-P300(CD) for H3K27ac deposition and transcription activation. **b**, Experimental timeline for ABA addition and cell collection at defined time points after ABA addition. **c**, Temporal enrichment of histone marks and transcriptional regulators at the IL1RN locus, including H3K27ac, H3K4me3, RNA pol-II Ser5Ph (initiation), NELF (pausing), p-TEFb (pause release), and RNA pol-II Ser2Ph (elongation), determined by ChIP-qPCR across multiple gene regions and time points. Fold changes were calculated relative to samples at t = 0 min (no-treatment). **d**, Curves of nonlinear regression analyses of ChIP enrichment for each mark and factor at distinct gene regions. t_50% maximum_ was calculated as the time to 50% maximum enrichment. Error bars represent mean ± s.e.m. from biological replicates (n = 3). Statistical significance was determined by two-way ANOVA. The detailed statistical analysis data of c and d including p-values and the R square for nonlinear regression can be found in Supplementary Table 1.

**Fig. 3.**
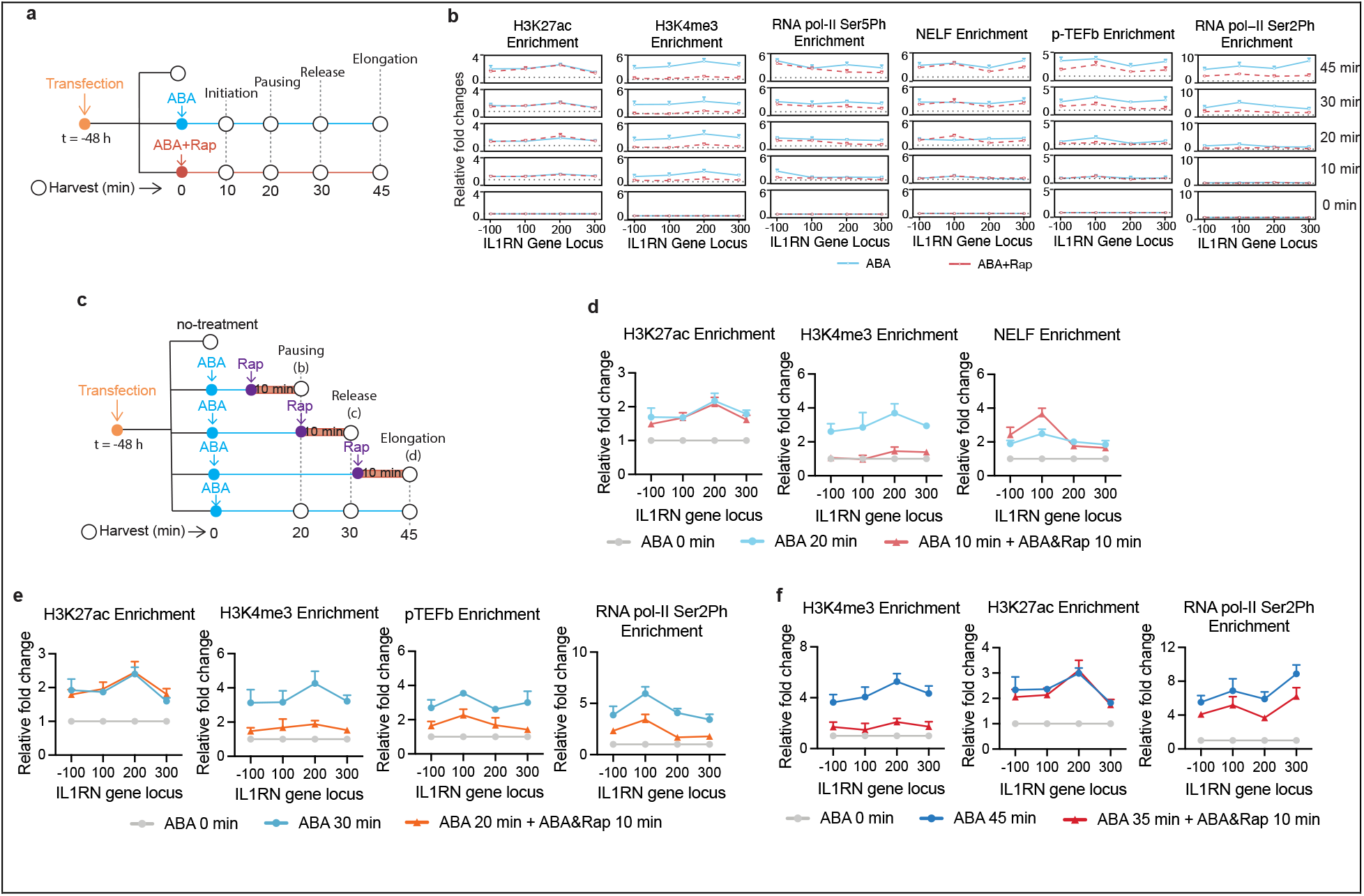
Stage-specific dissection of H3K27ac-induced transcription via acute H3K4me3. **a**, Time course experiments to determine H3K4me3 effects on different transcription stages by comparing conditions of H3K27ac-induced gene expression (+ABA) with or without H3K4me3 (- or + Rap). **b**, The enrichment of H3K27ac, H3K4me3, RNA pol-II Ser5Ph (initiation), NELF (pausing), p-TEFb (pause-release), and RNA pol-II Ser2Ph (elongation) at the IL1RN locus under ABA-only or ABA+Rap conditions at indicated time points, quantified by ChIP-qPCR assays. **c**, Time course experiments to study the impact on H3K27ac-induced gene expression by removing H3K4me3 (+Rap) at specific transcription stages after H3K27ac deposition (ABA). **d**, The enrichment of H3K4me3, H3K27ac and NELF (pausing factors) after cells were treated with ABA (writing H3K27ac) for 10 min, then with or without Rap (remove or maintain H3K4me3) for another 10 min before cells were collected and analyzed by ChIP-qPCR assays. **e**, The enrichment of H3K4me3, H3K27ac, p-TEFb (pause release) and RNA pol-II Ser2Ph (early elongation) after cells were treated with ABA for 20 min, then with or without Rap for another 10 min before cells were collected and analyzed by ChIP-qPCR assays. **f**, The enrichment of H3K4me3, H3K27ac and RNA pol-II Ser2Ph (elongation) after cells were treated with ABA for 35 min, then with or without Rap for another 10 min before cells were collected and analyzed by ChIP-qPCR assays. Fold changes were calculated relative to samples at t = 0 min. Error bars represent mean ± s.e.m. from biological replicates (n = 3). Statistical significance was determined by two-way ANOVA. The detailed statistical analysis data (p-values) can be found in Supplementary Table 1.

### The Effects of H3K4me3 on Initiation, Pausing and Elongation During Transcription

We next employed our dual editing system to investigate the stage-specific function of H3K4me3 at the IL1RN locus. To confirm that H3K4me3 can be erased rapidly, we determined the kinetics of Rap-induced recruitment of HA-FRB-KDM5B(CD) and H3K4me3 depletion under the background of H3K27ac-induced gene activation. Cells were co-transfected with Flag-ABI-P300(CD), HA-FRB-KDM5B(CD), PYL-dCas9-FKBP-V5 and IL1RN sgRNA for 48 h, followed by ABA treatment to induce H3K27ac/H3K4me3 deposition and transcription. 60 min after ABA addition, Rap was added to remove H3K4me3, and cells were harvested across a short time course (5, 15, 30 and 60 min after Rap addition; Extended Data Fig. 6a). ChIP-qPCR analyses revealed the rapid recruitment of HA-FRB-KDM5B(CD) and reduction of H3K4me3 within 5 min after Rap addition (Extended Data Fig. 6b), which supported the feasibility of using this tool to study transcription stage-specific roles of H3K4me3 within the expected transcription timeframe.

To examine how the H3K27ac-driven transcription process proceeds in the absence of H3K4me3, we first defined the precise time course of transcription stages following ABA addition including initiation (10 min), promoter-proximal pausing (20 min), pause-release (30 min), and elongation (45 min). These time points represent the approximate midpoint of each stage based on the determined t_50% maximum_ (Fig. 2d), ensuring that the chosen time points capture periods when each process is actively engaged. We used our dual editing tool to remove H3K4me3 from the very beginning during H3K27ac-driven transcription and observed the resulting impacts on each transcription stage. Cells were transfected with the dual editing modules and treated with either ABA or ABA+Rap for 10, 20, 30 and 45 min before harvesting and ChIP-qPCR analyses (Fig. 3a). In cells treated with ABA+Rap, H3K4me3 was efficiently depleted while H3K27ac remained intact (Fig. 3b). At the transcription initiation stage (10 min), RNA pol-II Ser5Ph was enriched at the TSS -100 bp site in cells treated with ABA alone (with intact H3K4me3), while in ABA+Rap treated cells (without H3K4me3), RNA pol-II Ser5Ph was not enriched at the same site until 20 min post-treatment, representing an approximate 10-min delay (Fig. 3b). At the promoter-proximal pausing stage, the timing of NELF recruitment at +100 bp was observed to be around 20 min in ABA+Rap treated cells, with no detectable delay compared to the ABA alone condition (Fig. 3b). However, the relative enrichment level of NELF in ABA+Rap treated cells was increased by approximately 30% at this position compared to cells with ABA alone, suggesting enhanced stabilization of promoter-proximal pausing in the absence of H3K4me3 (Fig. 3b). The recruitment of both p-TEFb (pause-release) and RNA pol-II Ser2Ph (elongation) was detected around 30 min at TSS +100 bp, and 45 min at +200 bp and +300 bp in ABA+Rap treated cells, representing a 10-min delay for pause-release and elongation compared to ABA alone treatment (Fig. 3b). Additionally, the overall enrichment levels of these elongation factors under the ABA+Rap condition were significantly decreased compared to cells treated with ABA alone, indicating that although transcription could progress to elongation without H3K4me3, it did so with reduced efficiency. These results suggest that H3K4me3 is required to achieve timely and efficient initiation, pause-release, and elongation, and its absence enhances promoter-proximal pausing, which stalls transcription.

It is possible that the observed effects at later transcription stages when lacking H3K4me3 were due to the cumulative effects from impaired earlier stages. To precisely dissect the impact of H3K4me3 on specific transcription stages, we performed stage-specific H3K4me3 removal by adding Rap 10 min prior to each defined stage time points. Cells were transfected as described above and treated first with ABA to induce the transcription, then with Rap at 10 min prior to indicated time points corresponding to different transcription stages before harvesting and ChIP-qPCR assays (treated at 10, 20, and 35 min and harvested at 20, 30, and 45 min) (Fig. 3c). Under these conditions, H3K27ac remained unaffected while H3K4me3 was efficiently erased at each stage (Fig. 3d-3f). Acute, stage-specific depletion of H3K4me3 led to a 35% increase in NELF recruitment at the TSS +100 bp site when compared to cell treated with ABA alone the entire period (p = 0.0005, Fig. 3d, Supplementary Table 1), indicating an enhanced stabilization of promoter-proximal pausing when losing H3K4me3. For pause-release and early elongation stages, we observed approximately 35% and 40% reductions in the recruitment of p-TEFb and RNA pol-II Ser2Ph, respectively, at the TSS +100 bp position (p-TEFb: p = 0.0062, RNA pol-II Ser2Ph: p = 0.0002, Fig. 3e, Supplementary Table 1). Productive elongation was also impaired when losing H3K4me3, with RNA pol-II Ser2Ph enrichment reduced by 40% at both +200 and +300 bp downstream of TSS (TSS +200bp: p = 0.0437, TSS +300bp: p = 0.018, Fig. 4f, Supplementary Table 1). Collectively, these results revealed critical and direct roles of H3K4me3 in pausing, pause-release, and elongation stages of transcription.

**Fig. 4.**
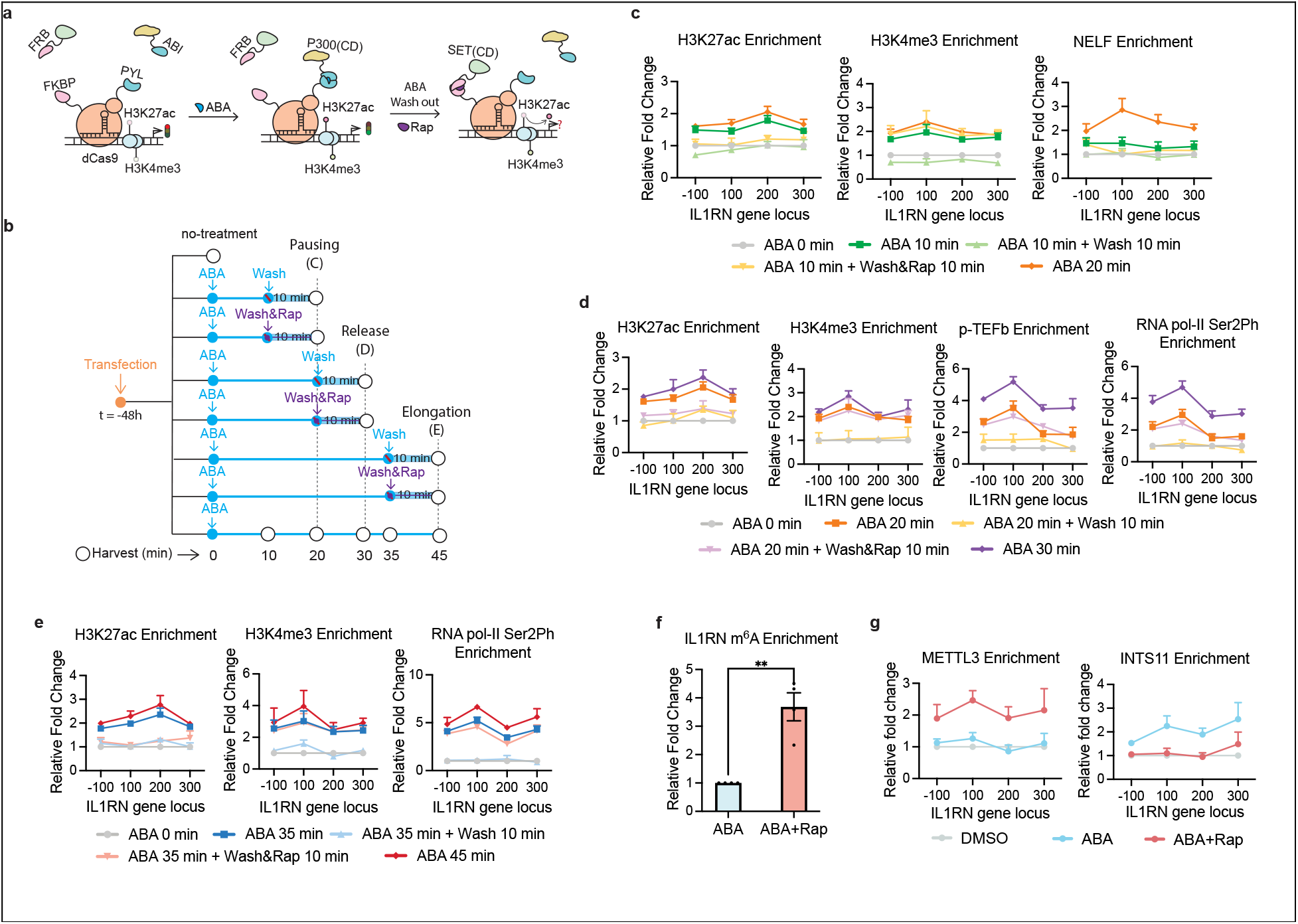
The impact of removing H3K27ac while maintaining H3K4me3 at specific transcription stages on transcriptional progression at the IL1RN locus. **a**, Using the dual-inducible system for temporal-specific manipulation of H3K27ac and H3K4me3 through adding or removing corresponding inducers. **b**, Time course experiments to study the impact of H3K27ac on each transcription stage and the role of H3Keme3 in the absence of H3K27ac in transcription. ABA was added to induce transcription, then removed with or without Rap added at 10 min before each stage to achieve acute H3K27ac depletion and H3K4me3 deposition. Cells were collected at 20 (pausing), 30 (pause release and early elongation), and 45 min (elongation). **c**, The enrichment of H3K27ac, H3K4me3, and NELF quantified by ChIP-qPCR assays from cells collected at 10 min before or at the pausing stage time point (20 min) with or without H3K27ac and/or H3K4me3. **d**, The enrichment of H3K27ac, H3K4me3, p-TEFb, and RNA pol-II Ser2Ph quantified by ChIP-qPCR assays from cells collected at 10 min before or at the pause release and early elongation stage time point (30 min) with or without H3K27ac and/or H3K4me3. **e**, The enrichment of H3K27ac, H3K4me3, and RNA pol-II Ser2Ph quantified by ChIP-qPCR assays from cells collected at 10 min before or at the elongation stage time point (45 min) with or without H3K27ac and/or H3K4me3. **f**, The enrichment of IL1RN mRNA with m^6^A by RIP, quantified by qPCR assays. Fold changes were calculated relative to cells treated with ABA alone. **g**, The enrichment of METTL3 and INTS11 quantified by ChIP-qPCR after ABA or ABA+Rap treatment. Fold changes were calculated relative to samples at t = 0 min (no-treatment) in (c-e) and to DMSO-treated cells in (f) and (g). Error bars represent mean ± s.e.m. from biological replicates (n=3 for c-e, n=4 for f and g). Statistical significance was assessed by two-way ANOVA for c-e and g and unpaired t-test for f. ns, P > 0.5; *P ≤ 0.05; **P ≤ 0.01; ***P ≤ 0.001; ****P ≤ 0.0001. The detailed statistical analysis data (p-values) can be found in Supplementary Table 1.

### Establishment of the H3K27ac and H3K4me3 Co-writing Platform

After investigating the direct effects of H3K4me3 on specific transcriptional stages, we investigated whether H3K27ac contributes to later stages in transcription beyond initiation. A key advantage of the ABA-inducible system is its reversibility,^30,37^ which allow us to rapidly remove H3K27ac at specific transcription stage and study the resulting impacts. To confirm the depletion of H3K27ac can occur in a timely fashion after ABA removal, we examined the kinetics regarding the dissociation of Flag-ABI-P300(CD) from the IL1RN locus following ABA washout, along with the subsequent changes of H3K27ac and H3K4me3 in the region (Extended Data Fig. 7a). We co-transfected cells with Flag-ABI-P300(CD), PYL-dCas9-FKBP-V5, and IL1RN sgRNAs for 48 h, then treated cells with ABA for 1 h to induce H3K27ac and H3K4me3 writing. ABA was then removed by washing cells with fresh medium, and cells were incubated for an additional 5, 15, 30 or 60 min without ABA before being harvested and analyzed by ChIP–qPCR assays (Extended Data Fig. 7b). We observed that Flag-ABI-P300(CD) dissociated from the IL1RN locus within 5 min after ABA washout (Extended Data Fig. 7c). Both H3K27ac and H3K4me3 levels decreased within 5 min and closing to the background level by 15 min (Extended Data Fig. 7c). These results demonstrated the highly dynamic nature of both modifications and the fast kinetics of our editing system for manipulating these histone marks.

After confirming the rapid reversal kinetics of H3K27ac and H3K4me3 established by the ABA-inducible editing system, we next sought to remove H3K27ac at defined transcription stages to investigate its roles beyond initiation. In addition, as opposed to our studies above to dissect the role of H3K4me3 in transcription with the presence of H3K27ac, we want to examine the impact of H3K4me3 in transcription when H3K27ac is absence. Since H3K4me3 quickly declined upon the depletion of H3K27ac (Extended Data Fig. 7c), to maintain H3K4me3 without H3K27ac, we engineered another dual editing system in which the Rap-inducible FRB–KDM5B(CD) eraser used in the studies above was replaced with a fusion of FRB to the H3K4me3-specific histone methyltransferase SET1A catalytic domain (SET(CD)) that allowed us to write H3K4me3 when H3K27ac disappeared (Extended Data Fig. 7d and 7e).^57^ To validate this system, cells were co-transfected with Flag-ABI-P300(CD), HA-FRB-SET(CD), PYL-dCas9-FKBP-V5 and IL1RN sgRNA and treated with ABA, Rap, ABA+Rap, or DMSO before analyzing by ChIP-qPCR. We confirmed that ABA treatment efficiently recruited Flag-ABI-P300(CD), resulting in increased H3K27ac, H3K4me3, and IL1RN expression (Extended Data Fig. 7f and 7g). Rap alone recruited HA-FRB-SET(CD), leading to elevated H3K4me3 but not H3K27ac, and failed to activate IL1RN expression (Extended Data Fig. 7f and 7g). Co-treatment with ABA and Rap efficiently recruited both Flag-ABI-P300(CD) and HA-FRB-SET(CD), leading to H3K27ac and H3K4me3 writing and IL1RN expression (Extended Data Fig. 7f and 7g).

We further validated the kinetics of HA-FRB-SET(CD) recruitment and H3K4me3 writing. We transfected cells were with HA-FRB-SET(CD), PYL-dCas9-FKBP-V5, and IL1RN-targeting sgRNAs and treated cells with Rap for 5, 15, 30 and 60 min before harvesting cells for ChIP-qPCR analyses (Extended Data Fig. 8a). We observed that both FRB–SET(CD) recruitment and H3K4me3 writing rapidly occurred within 5 min (Extended Data Fig. 8b), confirming the fast kinetics of the H3K4me3 writing platform.

### H3K27ac Depletion Collapses Transcription but Sustaining H3K4me3 Preserves Partial Transcriptional Output

We next leveraged this new dual editing system to stage-specifically remove H3K27ac while maintaining H3K4me3 at these transcription stages (Fig. 4a). To achieve complete H3K27ac depletion and H3K4me3 deposition in these stages, based on our kinetics studies (Extended Data Fig. 7c and 8b), ABA washout and Rap addition were performed 10 min in advance of each stage-representing time point (Fig. 4b). We transfected cells with Flag-ABI-P300(CD), HA-FRB-SET(CD), PYL-dCas9-FKBP-V5 and IL1RN sgRNA and treated cells with ABA, followed by ABA washout (removing H3K27ac) at the indicated time points, with or without Rap addition (writing H3K4me3) at the same time. Untreated cells were used as negative controls. We observed that when ABA was removed without adding Rap, both H3K27ac and H3K4me3, as well as all transcription stage markers, rapidly returned to baseline levels at all time points (the “ABA x min + Wash 10 min” condition in Fig. 4c-4e). By contrast, when ABA was removed but Rap was added, H3K27ac still reduced to background, but H3K4me3 levels remained significantly enriched (“ABA x min + Wash & Rap 10 min” in Fig. 4c-4e). Notably, under these conditions, NELF was reduced to background levels when H3K27ac was depleted even with H3K4me3 maintained, indicating that H3K27ac is necessary for sustaining the pausing stage, and H3K4me3 alone cannot induce or maintain promoter-proximal pausing (Fig. 4c). For pause-release and early elongation, both p-TEFb and RNA pol-II Ser2Ph were retained at levels comparable to the levels before ABA was removed (“ABA 20 min” in Fig. 4d), but less than the levels under the condition maintaining ABA the entire period (“ABA 30 min” in Fig. 4d), indicating that H3K4me3 can maintain the levels of p-TEFb and RNA pol-II Ser2Ph established by H3K27ac, but H3K4me3 alone cannot further facilitate transcription without H3K27ac. A similar trend was observed at the late elongation stage, where RNA pol-II Ser2P enrichment was preserved at the same level as before ABA washout (“ABA 35 min” in Fig. 4e) but did not reach the level observed when ABA remained for the entire time (“ABA 45 min” in Fig. 4e). These findings indicated that while H3K4me3 can maintain the transcription status established by H3K27ac, it cannot substitute H3K27ac for achieving optimal transcription progression and output. Collectively, these experiments demonstrate that H3K27ac is essential not only for transcription initiation but also for the effective progression of active transcription states.

### H3K4me3 Depletion Promotes mRNA Destabilization Through m^6^A Depositon

We observed that depleting H3K4me3 not only reduced the rate of transcription but also destabilized mRNA (Fig. 1i). Given that N6-methyladenosine (m^6^A) modification is a widely recognized mechanism for regulating mRNA stability,^58,59^ we examined whether the m^6^A level on IL1RN mRNA transcripts changed when H3K4me3 was depleted. We transfected cells with Flag-ABI-P300(CD), HA-FRB-KDM5B(CD), PYL-dCas9-FKBP-V5 and IL1RN-targeting sgRNAs, followed by treating cells with ABA or ABA+Rap. Total RNAs were harvested, and RNA immunoprecipitation (RIP) and RT-qPCR assays were performed to quantify the m^6^A level on IL1RN transcripts. We observed a significant increase in m^6^A on IL1RN mRNAs when H3K4me3 was depleted in the ABA+Rap treated cells compared to cells treated with ABA alone (Fig. 4f).

To investigate whether the increase in m^6^A deposition on IL1RN mRNAs resulted from the increased recruitment of m^6^A methyltransferase METTL3 to the IL1RN gene locus, we also performed ChIP-qPCR in the experiments above to quantify METTL3 occupancy at the IL1RN locus under both ABA and ABA+Rap conditions. We observed that the depletion of H3K4me3 led to a significant increase in METTL3 enrichment near the IL1RN TSS site (Fig. 4g), which was positively correlated with the elevated m^6^A level observed on the transcripts. It has been shown that METTL3 antagonizes the chromatin localization of integrator complex INTS11, suggesting potential competition between METTL3 and INTS11 for chromatin targeting.^60^ Moreover, H3K4me3 has been shown to be required for recruiting INTS11.^21^ Depleting H3K4me3 can potentially reduce INTS11 occupancy on chromatin that facilitated METTL3 localization. To examine this possibility, we performed ChIP-qPCR experiments as described above and observed that H3K4me3 depletion reduced INTS11 enrichment at the IL1RN TSS (Fig. 4g), suggesting the potential mechanistic connection between H3K4me3 depletion and METTL3 enrichment via INTS11. Collectively, our findings suggested a mechanistic interplay between H3K4me3 and mRNA stability and showed that H3K4me3 plays versatile roles in regulating gene expression beyond transcription.

## DISCUSSION

In this study, we developed a dual chemically inducible CRISPR-based epigenome editing system that enables rapid, reversible and independent manipulation of H3K4me3 and H3K27ac at specific gene loci. This system provides unique opportunities to temporal precisely edit chromatin states, offering insights into how specific histone modifications cooperatively shape transcriptional outcomes This platform enables transcription stage-specific modulation of H3K27ac and H3K4me3 at defined genomic loci with a minute-scale temporal resolution. Using this system, we systematically dissected the individual and cooperative contributions of H3K27ac and H3K4me3 to transcriptional activation and progression. Our results reveal that H3K27ac acts as an upstream transcriptional trigger and H3K4me3 is essential for supporting transcriptional output. Acute, stage-specific removal of H3K4me3 impairs RNA pol-II recruitment, increases promoter-proximal pausing and diminishes productive elongation. Surprisingly, losing H3K4me3 during transcription accelerates mRNA decay via enhanced (m^6^A) deposition, suggesting a regulatory role of H3K4me3 in post-transcriptional gene regulation. Conversely, the loss of H3K27ac, resulting in subsequent loss of H3K4me3, rapidly abolishes transcriptional activity, while the maintenance of H3K4me3 without H3K27ac after transcription initiation preserves partial capacity to recruit transcriptional machinery but cannot substitute H3K27ac for achieving the full extent of robust transcription. In conclusion, our studies elucidate the functional hierarchy and interdependence of H3K27ac and H3K4me3 during dynamic transcriptional regulation. The dual-inducible epigenome editing system provides a powerful and modular platform that can be modified and applied to study the functional relationship between histone modifications during gene expression. Future studies incorporating additional epigenetic and transcriptional regulators will further illuminate how chromatin modifications orchestrate dynamic gene regulation in cellular processes.

## METHODs

### Plasmid Construction

All DNA fragments were amplified by polymerase chain reaction (PCR) from intermediate constructs using CloneAmp HiFi PCR Premix (Takara #639298) on an S1000 thermal cycler (Bio-Rad). Primer sequences used to generate individual constructs for dCas9, effector domains, and sgRNA nucleotide sequences are provided in the Extended Data Table 1 and 2.

Human codon-optimized Streptococcus pyogenes dCas9 containing three C-terminal SV40 nuclear localization signals (NLSs) was derived from a previously reported dCas9-PYL-HA construct and further engineered to incorporate a synthesized 3×FKBP sequence. For transcriptional effector fusions, Flag-ABI-P300(CD) (amino acids 1048–1664) was cloned and codon-optimized based on a previously reported Flag-ABI-P300(CD), with a flexible XTEN linker inserted between ABI and P300(CD). For rapamycin-inducible recruitment, the H3K4me3/2-specific demethylase KDM5B catalytic core domain (amino acids 3–846) was cloned from Addgene plasmid #143615 (a gift from Feng Zhang). The SET catalytic domain of SET1A (amino acids 1414–1707) was cloned from a previously reported dCas9-SET(CD) fusion.

### Cell culture and transfection

HEK293T cells were cultured in Dulbecco’s modified Eagle’s medium (DMEM; Gibco #11995073) supplemented with 10% fetal bovine serum (FBS; Biotechne|R&D systems #S11550), 1% penicillin–streptomycin (Gibco #15140122), and 1% GlutaMAX (Gibco #35050061) in a humidified incubator at 37 °C with 5% CO_2_. Cells were seeded at a density of 4 × 10^5^ cells per ml in either 24-well plates or 10-cm dishes one day before transfection.

For transfection with the dual editing modules, 10-cm dishes were transfected with 4 µg PYL-dCas9-FKBP, 6 µg Flag-ABI-P300(CD), 6 µg HA-FRB-KDM5B(CD) or HA-FRB-SET(CD), and a total of 1 µg sgRNAs (250 ng each). Plasmid DNA was mixed with 36 µL P3000 reagent (Thermo Fisher #L3000015) in 500 µL Opti-MEM (Gibco #31985062), and in parallel, 36 µL Lipofectamine 3000 (Thermo Fisher #L3000015) was diluted in 500 µL Opti-MEM. After 5 min, the two mixtures were combined and incubated for 15 min at room temperature before being added to the cells. Transfected cells were cultured for an additional 24–48 h prior to downstream assays. For 24-well plate transfections, 50 µL of the prepared transfection mixture was added per well.

### Expression of proteins

Twenty-four hours before transfection, HEK293T cells were seeded in 6-well plates at a density of 4 × 10^5^ cells per mL. Cells were transfected with either 800 ng of dCas9 fusion constructs or 1.2 µg of Flag–ABI–P300 or HA–FRB–KDM5B(CD)/HA–FRB–SET(CD) using Lipofectamine 3000 (Thermo Fisher) according to the manufacturer’s instructions.

At 24 h post-transfection, cells were washed twice with 2 mL ice-cold PBS and lysed on ice with 100 µL RIPA buffer (Thermo Fisher #89901) supplemented with a protease inhibitor cocktail (Thermo Fisher #89901). Lysates were gently vortexed and incubated on ice for 30 min, followed by centrifugation at 16,000 × g for 20 min at 4 °C. Supernatants were collected, and protein concentrations were determined using the Pierce 660 nm Protein Assay (Thermo Fisher #22660) according to the manufacturer’s instructions.

Equal amounts of protein (20 µg per sample) were separated by SDS–PAGE at 120 V for 90 min and transferred onto PVDF membranes (Biorad #1620177) at 10 mA for 16 h at 4 °C. Membranes were blocked for 1 h at room temperature in 5% non-fat milk in TBST, washed three times with TBST, and incubated for 2 h at room temperature with primary antibodies against HA (CST#3724S), Flag (CST#14793S), V5 (CST#13202S), or GAPDH (#5174S). After three washes with TBST, membranes were incubated with the HRP-conjugated anti-Rabbit IgG secondary antibodies (CST #7074) for 1 h, washed again, and visualized using the ChemiDoc™ MP Imaging System (Bio-Rad). GAPDH served as a loading control.

### Chromatin immunoprecipitation (ChIP) assay

HEK293T cells were seeded in 10-cm or 5-cm dishes at a density of 4 × 10^5^ cells per mL in culture medium 24 h prior to transfection. Cells were transfected with 4 µg PYL-dCas9-FKBP, 6 µg Flga-ABI-P300(CD), 6 µg HA-FRB–X [KDM5B(CD) or SET(CD)], and 2 µg sgRNA plasmid using Lipofectamine 3000 according to the manufacturer’s protocol.

At 24 or 48 h post-transfection, cells were treated with 100 µM abscisic acid (ABA; Glod Biotech #A-050-5), 20 nM rapamycin (Rap; SantaCruz#sc-3504), or an equal volume of DMSO as control, and incubated for 24 h or for the indicated shorter durations depending on the experimental design. At the designated time point, cells were crosslinked with 1% formaldehyde (Sigma Aldrich#F8775) at room temperature for 10 min. Crosslinking was quenched by adding 1 mL of 1 M glycine (FisherScinetic#AAJ16440736) to each 10 mL of culture medium and incubating for 5 min at room temperature. Media were removed, and cells were washed twice with 10 mL of ice-cold PBS.

Cells were scraped in 1 mL ice-cold PBS supplemented with protease inhibitor cocktail (PIC; ThermoFisher#78438) and pelleted by centrifugation at 1,000 × g for 5 min at 4 °C. The pellets were resuspended in cell lysis buffer, incubated on ice for 10 min, and centrifuged at 5,000 × g for 5 min at 4 °C; this step was repeated twice. Nuclei were lysed in nuclear lysis buffer for 10 min on ice, and chromatin was fragmented by sonication at 4 °C using a Bioruptor Pico (Diagenode) for 10 cycles of 30 son/30 s off.

For each ChIP reaction, 100 µL of sheared chromatin was combined with 410 µL 1× ChIP buffer (CST#7008), the indicated ChIP-grade antibody (anti-H3K27ac CST#8173S; anti-H3K4me3 CST#9751S; Anti-Flag CST#14793S; Anti-HA CST#3724S; Anti-RNA pol II CTD Ser 5Ph CST#13499S; Anti-TH1L CST#12265S; Anti-CyclinT1 CST#81464S; Anti-RNA pol II CTD Ser2Ph CST#13523S; Anti-METTL3 CST#86132S), and PIC. 2% of each sample was saved as input control. Reactions were rotated overnight at 4 °C, followed by incubation with 30 µL ChIP-grade Protein G magnetic beads (CST#9006S) for 2 h at 4 °C. Beads were sequentially washed three times with 1 mL low-salt wash buffer (1× ChIP buffer) and once with 1 mL high-salt wash buffer (1× ChIP buffer supplemented with 70 µL of 5 M NaCl). Bound chromatin was eluted with 150 µL ChIP Elution Buffer at 65 °C for 30 min. Crosslinks were reversed by incubation at 65 °C for ≥2 h in the presence of proteinase K (Thermo Scientific #EO0491) and 6 µL of 5 M NaCl. DNA was purified using the ChIP DNA Purification Kit (CST#14209) and analyzed by qPCR.

### Total RNA extraction and purification

Total RNA was extracted from HEK293T cells using TRIzol reagent (Thermo Fisher#15596026) according to the manufacturer’s instructions, with chloroform (Fisher Scientific#AAJ67241), isopropanol (Fisher Scientific#AC317270010), and 75% ethanol for phase separation and precipitation. Briefly, HEK293T cells were seeded in 24-well plates or 10-cm dishes at a density of 4 × 10^5^ cells/mL 24 h prior to transfection. At 24 h post-transfection, cells were treated with 100 µM abscisic acid ABA (Glod Biotechnology #A-050-5), 20 nM Rap (Santa Cruz #sc-3504), or an equivalent volume of DMSO (negative control) and incubated for an additional 24 h. For large-scale preparations, cells cultured in 10-cm dishes were lysed directly with 2 mL TRIzol, collected by pipetting, and incubated on ice for 3–5 min. Chloroform (0.2 mL per 1 mL TRIzol) was added, samples were shaken vigorously for 2 min, incubated on ice for 10–15 min, and centrifuged at 12,000 × g for 15 min at 4 °C. The aqueous phase was transferred to a fresh tube, and RNA was precipitated with isopropanol (0.5 mL per 1 mL TRIzol), incubated on ice for 10– 30 min, and centrifuged at 12,000 × g for 10 min at 4 °C. Pellets were washed once with 75% ethanol, air dried, and resuspended in 50–100 μL nuclease-free water. Purified RNA was either immediately subjected to reverse transcription or stored at −80 °C.

### RNA immunoprecipitation

Total RNA was extracted from cells cultured in 6-well plates using TRIzol reagent. For each reaction, 10 µg of total RNA was incubated with 3 µL of N6-methyladenosine (m6A) antibody (CST, #56593) in 250 µL of Reaction Buffer (150 mM NaCl, 10 mM Tris-HCl, pH 7.5, 0.1% NP-40 in nuclease-free H2O) supplemented with 3 µL of RNasin® Plus inhibitor (Promega, #N2611) for 1 h at 4 °C with orbital rotation. Protein G magnetic beads (25 µL per reaction; NEB, #S1430) were prewashed twice with 200 µL of reaction buffer and subsequently added to the RNA– antibody mixture, followed by incubation for 2 h at 4 °C with rotation. The beads were sequentially washed twice with 200 µL of Reaction Buffer, twice with 200 µL of Low-Salt Buffer (50 mM NaCl, 10 mM Tris-HCl, pH 7.5, 0.1% NP-40), and twice with 200 µL of High-Salt Buffer (500 mM NaCl, 10 mM Tris-HCl, pH 7.5, 0.1% NP-40). Bound RNA was eluted with 30 µL of Buffer RLT (Qiagen, #79216) for 1 min at room temperature.

To purify the RNA, 20 µL of Dynabeads MyOne Silane (Life Technologies, #37002D) prewashed with 100 µL of Buffer RLT and resuspended in 30 µL of Buffer RLT were added to the eluate. Subsequently, 60 µL of absolute ethanol was mixed with the RNA–Dynabeads suspension and incubated for 1 min at room temperature. The beads were washed twice with 200 µL of 70% ethanol and air-dried for 5 min at room temperature. Purified RNA was eluted with 20 µL of RNase-free water (Gibco) by incubation for 1 min at room temperature. The recovered RNA, together with 1 µg of input total RNA, was used for cDNA synthesis and RT-qPCR analysis.

### Reverse transcription

Complementary DNA (cDNA) was synthesized from 200 ng of total RNA using the iScript cDNA Synthesis Kit (Bio-Rad#1708890) according to the manufacturer’s instructions. Each 20 μL reaction contained 4 μL 5× reaction mix, 1 μL reverse transcriptase, nuclease-free water, and RNA template. Reverse transcription was performed in a thermal cycler (Bio-Rad) under the following program: 25 °C for 5 min (priming), 46 °C for 25 min (reverse transcription), and 95 °C for 1 min (enzyme inactivation). The resulting cDNA was either used immediately for qPCR analysis or stored at −20 °C.

### Quantitative q-PCR

For both ChIP-derived DNA and reverse-transcribed cDNA, 1 μL of DNA template was used per reaction. qPCR was performed using SsoAdvanced Universal SYBR Green Supermix (Bio-Rad#1725274) in a 10 μL total volume containing 5 μL 2× SYBR Green mix, 2 μL primer mix (0.25 μM each forward and reverse), nuclease-free water, and template DNA, according to the manufacturer’s instructions. Reactions were conducted in a 384-well format on a 7900HT Fast Real-Time PCR System (Applied Biosystems), with all assays performed in at least technical triplicates. Raw data were collected and analyzed using SDS 2.4 software under default parameters.

For gene expression analysis, GAPDH served as the internal control. For ChIP–qPCR, enrichment was normalized to 2% input DNA. Thermal cycling conditions were as follows:

#### Polymerase activation/initial denaturation

95 °C for 30 s (cDNA) or 95 °C for 3 min (ChIP DNA)

#### Denaturation

95 °C (cDNA) or 98 °C (ChIP DNA) for 15 s

#### Annealing/extension

60 °C for 1 min with plate read, repeated for 40 cycles

#### Dissociation curve

95 °C for 15 s, 60 °C for 1 min, 95 °C for 15 s (See also Extended Data Table 3 for q-PCR primer sequence)

### Quantification and statistical analysis

All statistical analyses were performed using GraphPad Prism 10 with default settings. Differences between experimental and control groups in ChIP-qPCR and gene expression assays were evaluated by one-way, two-way ANOVA and unpaired t-test, as appropriate. In this manuscript, a p value less than 0.05 was marked as *, less than 0.01 as ** and less than 0.001 as ***. Detailed p values and R^2^ values from nonlinear regression analyses are provided in Supplementary Table 1.

## Data and code availability

Any additional information required to reanalyze the data reported in this paper is available from the lead contact upon request.

## ACKNOWLEDGMENTS

This work was supported by NIH R01GM143256.

## AUTHOR CONTRIBUTIONS

F-S.L. and C.Z. conceived the project and designed the experiments unless otherwise stated. C.Z. performed most of the experiments, analyzed data and prepared all the Fig.s. C.D. performed RNA immunoprecipitation (RIP) – qPCR. W.Z. contributed to plasmids construction. F-S. L. and C.Z. wrote the manuscript.

