## Supplementary information for "Hierarchical Interplay between H3K27ac and H3K4me3 in Transcriptional Regulation"

**This PDF file include Extended Data Fig 1-8 and Extended Data Table 1-3.**

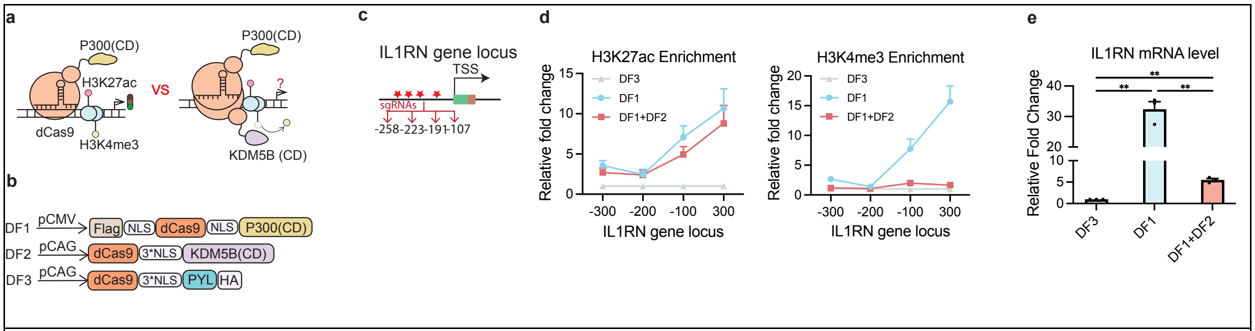


**Extended Data Fig 1 | dCas9-P300/KDM5B-mediated editing of H3K27ac and H3K4me3.** **a,** dCas9 fusion proteins-based epigenome editing. Left: dCas9-P300(CD) for H3K27ac writing and gene expression; right: dCas9-P300(CD) and dCas9-KDM5B(CD) for both H3K27ac writing and H3K4me3 erasing. **b,** dCas9 fusion protein constructs for epigenome editing. **c,** The IL1RN promoter and sgRNA target sites (red stars and the corresponding sites relative to TSS). **d,** ChIP-qPCR analysis of H3K27ac (left) and H3K4me3 (right) enrichment at the IL1RN promoter (-300 bp, -200 bp, -100 bp and 300 bp from the TSS) following recruitment of dCas9-P300(CD) (DF1), dCas9–PYL (DF3), or both dCas9–P300(CD) and dCas9–KDM5B(CD) (DF1+DF2). **e,** IL1RN mRNA levels measured by RT-qPCR under the indicated treatments. Data represent mean ± s.e.m. (n = 3). Statistical significance was determined by two-way ANOVA (d) or one-way ANOVA (e). ns, $P > 0.5$; *$P \leq 0.05$; **$P \leq 0.01$; ***$P \leq 0.001$; ****$P \leq 0.0001$. The detailed statistical analysis data (p-values) can be found in Supplementary Table ***1.***

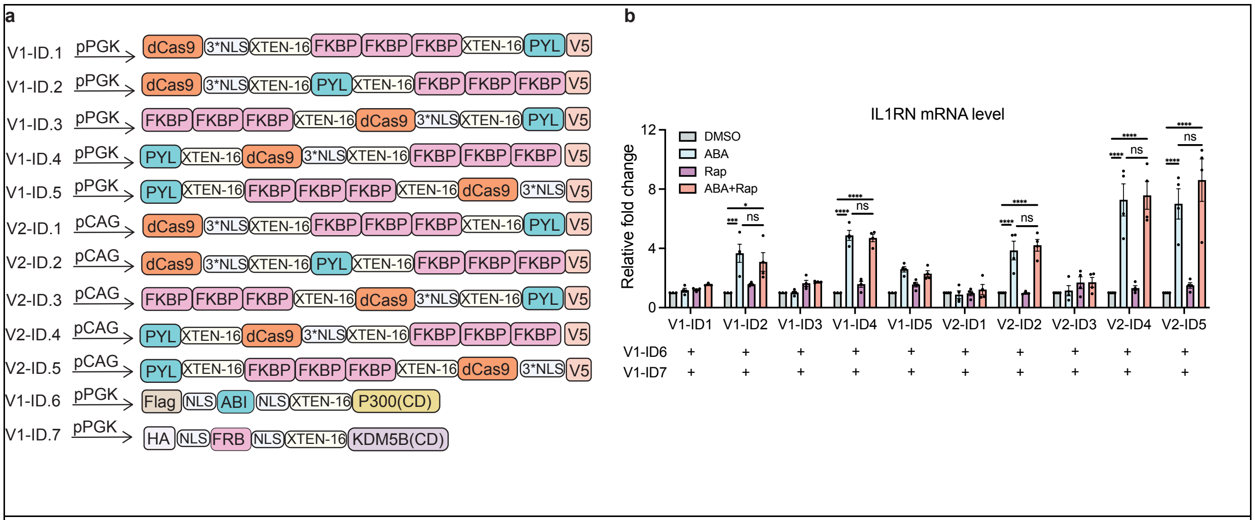


**Extended Data Fig 2 | The screening and optimization of dCas9-based dual epigenome editing system. a,** Constructs of PYL/dCas9/3XFKBP, fusion protein with low or high promoters pPGK and pCAG (V1/2-ID1-5); the construct scheme of pPGK-Flag-ABI-P300(CD) (V1-ID6); the construct scheme of pPGK-HA-FRB-KDM5B(CD) (V1-ID7). **b,** IL1RN mRNA levels quantified by RT-qPCR in cells expressing the indicated combinations of V1/2-ID1-5 with V1-ID6 and V1-ID7 after treatment with DMSO, ABA, Rap, or ABA+Rap. The enrichment fold changes were calculated by comparing to the DMSO-treated samples. Error bars represent mean ± s.e.m. from biological replicates (n = 4). Statistical significance was determined by two-way ANOVA. ns, $P > 0.5$; * $P \leq 0.05$; ** $P \leq 0.01$; *** $P \leq 0.001$; **** $P \leq 0.0001$. The detailed statistical analysis data (p-values) can be found in Supplementary Table ***1***

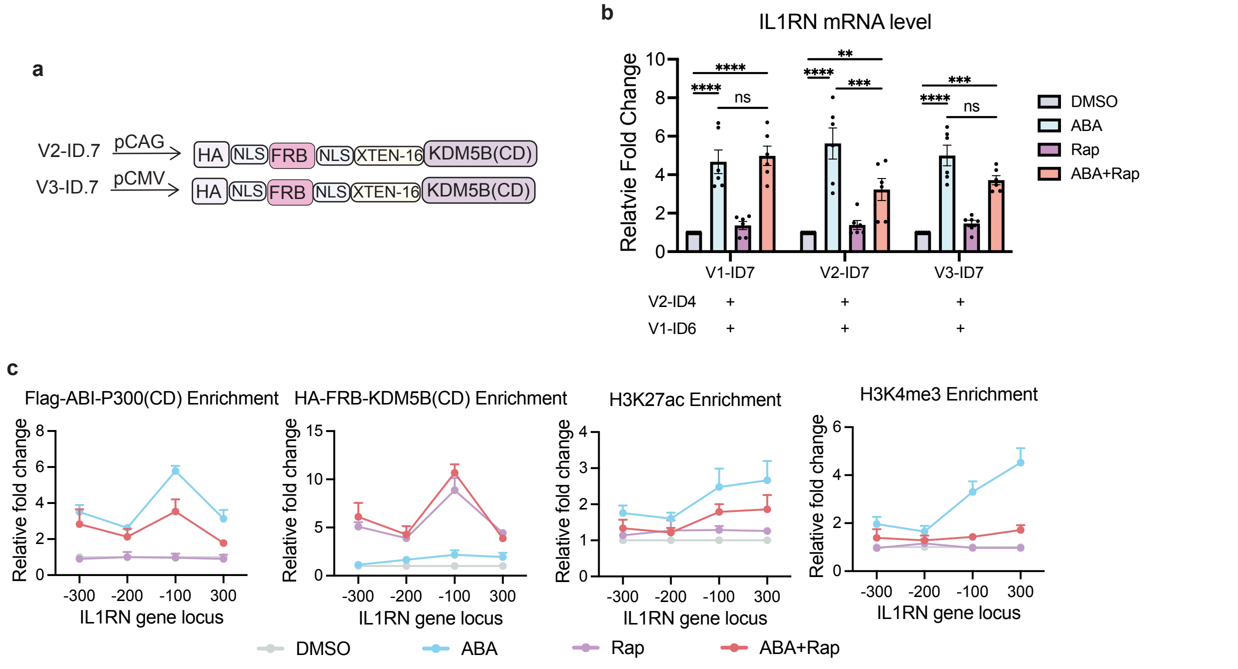


**Extended Data Fig 3 | Promoter Testing and Functional Validation of dCas9-Based Dual Epigenome Editing Configurations a,** Constructs of HA-FRB-KDM5B(CD) driven by pCAG (V2) or pCMV (V3). **b,** IL1RN mRNA levels following co-expression of V2-ID4, V1-ID6, and V1, V2 or V3-ID7. **c,** Enrichment of Flag-ABI-P300(CD), HA-FRB-KDM5B(CD), H3K27ac, and H3K4me3 at the IL1RN promoter quantified by ChIP-qPCR assays following DMSO, ABA, Rap, or ABA+RAP treatment. The enrichment fold changes were calculated by comparing to the DMSO-treated samples. Error bars represent mean ± s.e.m. from biological replicates (n=6 for b, n = 4 for c). Statistical significance was determined by two-way ANOVA. ns, $P > 0.5$; * $P \leq 0.05$; ** $P \leq 0.01$; *** $P \leq 0.001$; **** $P \leq 0.0001$. The detailed statistical analysis data (p-values) can be found in Supplementary Table ***1***

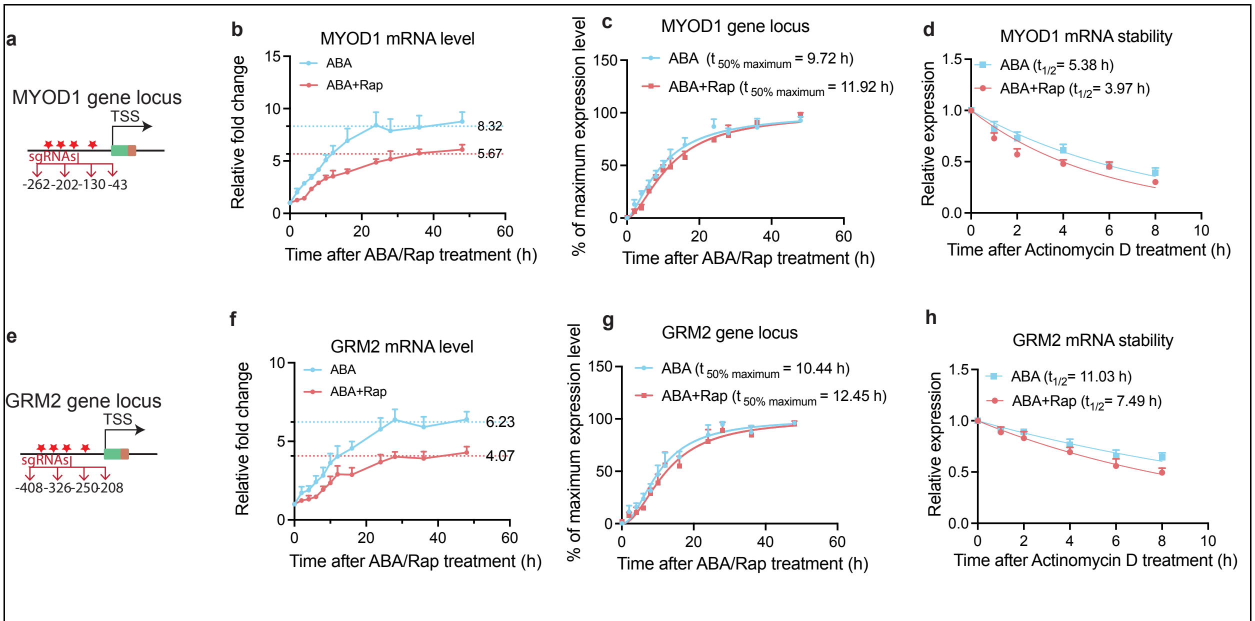


**Extended Data Fig 4 | H3K4me3 regulates mRNA production and stability at MYOD1 and GRM2 gene loci. a,** The MYOD1 promoter and sgRNA target sites (red stars and the corresponding sites relative to TSS). **b,** Time-course of MYOD1 mRNA accumulation measured by RT-qPCR after ABA (steady-state mRNA level: 8.32) or ABA+Rap treatment (steady-states mRNA level: 5.67). **c,** Nonlinear regression analysis of MYOD1 expression rate. $t_{50\%\ maximum}$ is defined as time reaches 50% maximum expression level. ($R^2$: 0.90 for ABA treatment; 0.92 for ABA+Rap treatment). **d,** mRNA decay curves of MYOD1 after actinomycin D treatment. $t_{1/2}$ is defined as the mRNA half-life ($R^2$: 0.84 for ABA treatment; 0.78 for ABA+Rap treatment). **e,** The GRM2 promoter and sgRNA target sites (red stars and the corresponding number). **f,** Time-course of MYOD1 mRNA accumulation measured by RT-qPCR after ABA (steady-state mRNA level: 6.23) or ABA+Rap treatment (steady-states mRNA level: 4.07). **g,** Nonlinear regression analysis of GRM2 expression rate ($R^2$: 0.90 for ABA treatment; 0.86 for ABA+Rap treatment). **h,** mRNA decay curves of GRM2 after actinomycin D treatment ($R^2$: 0.75 for ABA treatment; 0.80 for ABA+Rap treatment). Fold changes calculated by comparing to samples at (t = 0 h) in (b) and (g). Fold changes were calculated relative to the expression level under ABA or ABA+Rap condition before Actinomycin D treatment (t = 0 h) in (d) and (h). Error bars represent mean ± s.e.m. from biological replicates (n = 4).

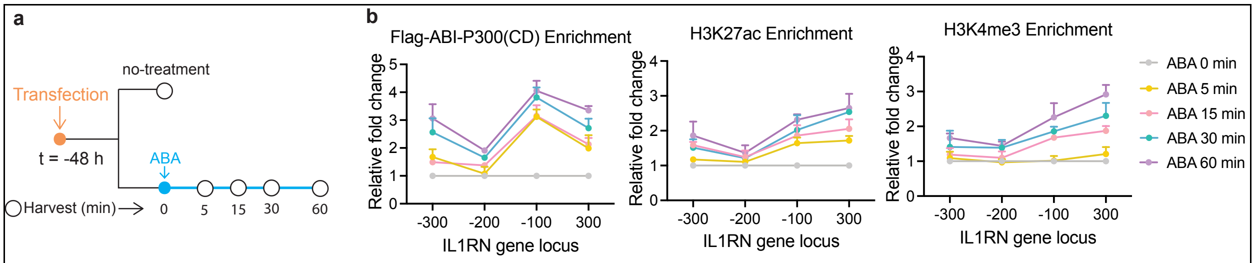


**Extended Data Fig 5 | Kinetics of chemically induced recruitment of Flag-ABI-P300(CD) and resulting H3K27ac and H3K4me3 writing. a,** Experimental timeline for ABA addition and cell harvesting at indicated time points after ABA addition. **b,** Flag-ABI-P300(CD) (left), H3K27ac (middle), and H3K4me3 (right) enrichments at the IL1RN locus following ABA treatment for 0, 5, 15, 30, and 60 min by ChIP-qPCR analyses. Fold changes were calculated relative to samples collected before ABA addition (no-treatment; t = 0 min). Error bars represent mean ± s.e.m. from biological replicates (n=3). Statistical significance was assessed by two-way ANOVA. The detailed statistical analysis data (p-values) can be found in Supplementary Table ***1***

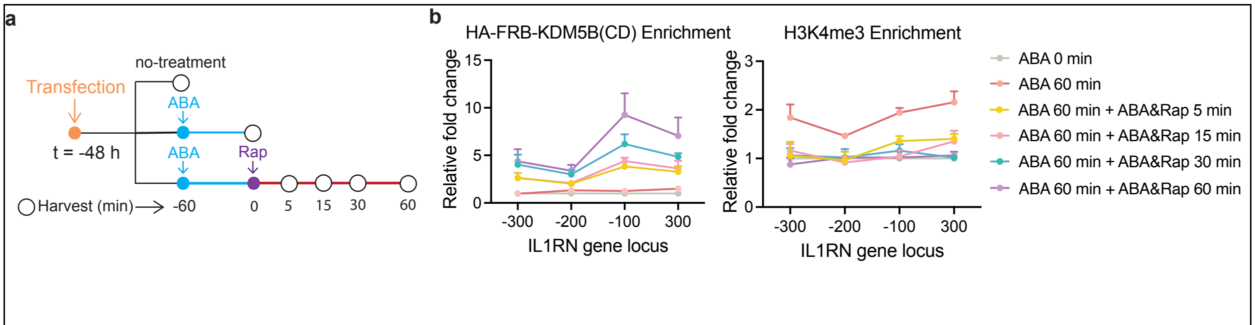


**Extended Data Fig 6 | Kinetics of chemically induced recruitment of HA-FRB-KDM5B(CD) and resulting H3K4me3 erasing. a,** Experimental timeline for ABA and rapamycin addition and cell harvesting at indicated time points after Rap addition to test the kinetics of H3K4me3 erasing. **b,** The enrichment of HA-FRB-KDM5B(CD) and H3K4me3 at the IL1RN locus following ABA and rapamycin treatment for different time periods, analyzed by ChIP-qPCR analyses. Fold changes were calculated relative to samples collected before ABA addition (no-treatment, t = -60 min). Error bars represent mean ± s.e.m. from biological replicates (n=3). Statistical significance was assessed by two-way ANOVA. The detailed statistical analysis data (p-values) can be found in Supplementary Table ***1***

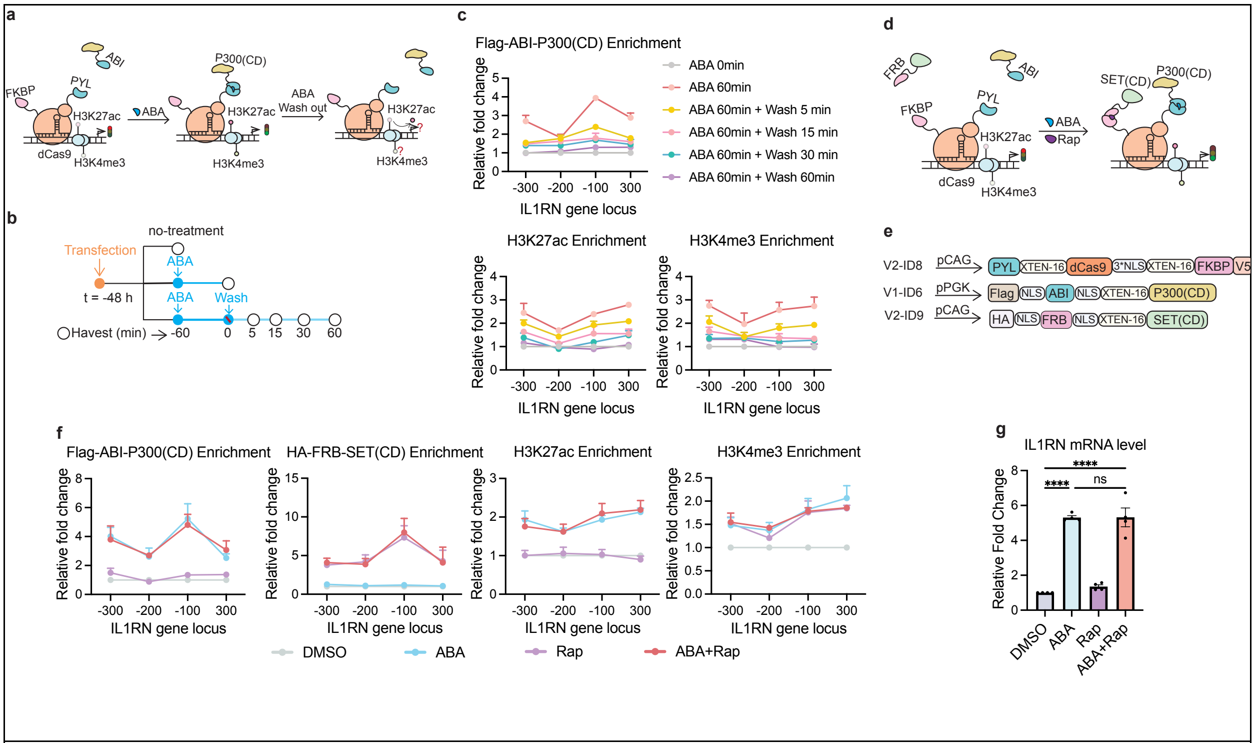


**Extended Data Fig 7 | Kinetics and reversibility of installed H3K27ac and H3K4me3, and the inducible H3K4me3 writing system by recruiting SET(CD) at the IL1RN locus**. **a,** Reversing ABA-induced H3K27ac and H3K4me3 deposition by the removal of ABA. **b,** Time course experiments to study the stability of ABA-induced deposition of H3K27ac and accompanied H3K4me3 after ABA removal. Transfected cells were treated with ABA for 60 min, followed by ABA removal and incubated for another 5, 15, 30 or 60 min before analyzed by ChIP-qPCR assays. **c,** The enrichment of Flag-ABI-P300(CD), H3K27ac, and H3K4me3 at indicated time points after ABA removal, quantified by ChIP-qPCR assays. **d,** The inducible H3K27ac and H3K4me3 dual writing system using HA-FRB-SET(CD) and Flag-ABI-P300(CD). **e,** Constructs of the CRISPR-based dual H3K27ac and H3K4me3 writing system. **f,** The enrichment of Flag-ABI-P300(CD), HA-FRB-SET(CD), H3K27ac, and H3K4me3 under DMSO, ABA, Rap, or ABA+Rap conditions, quantified by ChIP-qPCR assays. **g,** IL1RN mRNA levels quantified by RT-qPCR under indicated conditions. Fold changes were calculated comparing to non-ABA treated cells (t = -60 min) in (b) and DMSO-treated samples in (e) and (f). Error bars represent mean ± s.e.m. from biological replicates (n = 3 for c and f, n=4 for g). Statistical significance was assessed by two-way ANOVA for c,f and one-way ANOVA for g. ns, $P > 0.5$; $*P \leq 0.05$; $**P \leq 0.01$; $***P \leq 0.001$; $****P \leq 0.0001$. The detailed statistical analysis data (p-values) can be found in Supplementary Table ***1***

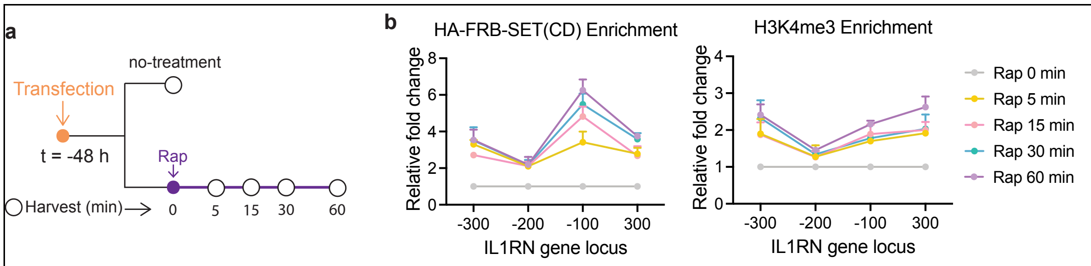


**Extended Data Fig 8 | Kinetics of chemically induced recruitment of HA-FRB-SET(CD) and resulting H3K4me3 writing. a,** Experimental timeline for Rap-induced recruitment of HA–FRB–SET(CD) and H3K4me3 writing. **b,** The enrichment of HA–FRB–SET(CD) and H3K4me3 at 0, 5, 15, 30, and 60 min after Rap treatment. Fold changes were calculated relative to samples collected before Rap addition (no-treatment; t = 0 min). Error bars represent mean ± s.e.m. from biological replicates (n=3). Statistical significance was assessed by two-way ANOVA. The detailed statistical analysis data (p-values) can be found in Supplementary Table ***1.***

### Extended Data Table 1 | Primers for PCR

| Plasmid | Amplicon | Fw (5'-3') | Rv (5'-3') |
|---|---|---|---|
| DF 2 | KDM5B(CD) | GGCAGCAGCGGGGGGTCAGCTAGCATGGAGGCGGCCACCACA | TGATCTAGAGTTCACGCGGCCGCACCGGTCTGCCGGAGCTCATTCACTGT |
| V1-ID1-5 | PGK promoter | CCACGCGGATCTCGGGTAGGGGAGGCGCTTTTC | ACGTTGCCCAGGAGCGAAAGGCCCGGAGATGAGG |
| V2-ID1 | XTEN-3XFKBP-XTEN | AGGAGGCTCCTCTGGCGGGTCAGGCGCGCCAACTCAAGAC | TACCGGGGCGATCGCATCCGGAGGATCTAGCGGAGG |
| V2-ID2 | XTEN-PYL | GGTAAATTCTACCGGGGCGATCGCATCCGGAGGTTCATCTGGC | AGATCCTCCGGATCCTGCAGGGTTCATAGCTTCAGTGATCGAAG |
| | XTEN-3XFKBP | CTGCAGGATCCGGAGGATCTAGCGGAGG | TAGGGATAGGCTTACCCCTAGGCTCCAGCTTGAGCAGCTCTAC |
| V2-ID3 | XTEN-3XFKBP | TCGAGCTCAAGCTTGCGGGACTGCAGATGAGCGGAGTGCAGGTGGAAACCATCTC | ATAGAATACTTCTTGTCCATCTGCAGATGCATTGACCCGCCAGAGGAGCCTCCT |
| V2-ID4 | PYL-XTEN | ATTTTGGCAAAGAATTCCTCGAGCTCATGACTCAAGACGAATTCACCCAA | CCGCAAGCTTTGACCCCCCGCTGCTGCC |
| | dCas9-XTEN | CGGGGGGTCAAAGCTTGCGGGACTGCAGAT | TTCCACCTGCACTCCGCTAGCTGACCCGCCAGAGGAGCCTCCT |
| V2-ID5 | XTEN-3XFKBP-XTEN | TTCGATCACTGAAGCTATGAACGCGATCGCATCCGGAGGATC | TTGTCCATCTGCAGTCCCGCTGGCGCGCCTGACCCGCCAGA |
| | dCas9-3*NLS | GCGGGACTGCAGATGGACAA | TTAGGGATAGGCTTACCCCTAGGCGCCCCGGTAGAATTTACCTT |
| V2-ID8 | 1XFKBP | CTACCCGAGATCCGCGTGGCCGTAC | GCCTTTCGCTCCTGGGCAACGTGCTGGT |
| V1-ID6 | PGK promoter | GCCAGATATACGCGGGTAGGGGAGGCGCTTTTC | CTCTAGTTAGCCAGACGAAAGGCCCGGAGATGAGG |
| V1-ID7 | PGK promoter | GCCAGATATACGCGGGTAGGGGAGGCGCTTTTC | CTCTAGTTAGCCAGACGAAAGGCCCGGAGATGAGG |
| V2-ID7 | HA-NLS-FRB-NLS | AGCTCAAGCTTGCGGGACTGCAGAGCGCCATGTACCCGTACGA | CTAGATCCTCCGGATGCGATCGCGCGCCCAACTTTGCGTTTC |
| V3-ID7 | HA-NLS-FRB-NLS | CTGGCTAGCGCCATGTACCCGTACGA | AGCGGTTTAAACTCAACCGGTCTGCCGGAGCTC |
| V2-ID9 | HA-NLS-FRB-NLS | CGAGCTCAAGCTTGCGGGAC | GGCTCGTAGGCATGCATTGACCCCCCGCTGCTGCCC |
| | SET (CD) | CCGGTGCGGCCGCGTGAA | GTCCCGCAAGCTTGAGCTCG |

**Extended Data Table 2 | sgRNA sequence[1–3]**

| Gene | Sequence | PAM | assembly |
|---|---|---|---|
| IL1RN promoter 1(+87 bp) | TTGTACTCTCTGAGGTGCTC | TGG | GRCh38.p14 |
| IL1RN promoter 2 (+238 bp) | TACGCAGATAAGAACCAGTT | TGG | GRCh38.p14 |
| IL1RN promoter 3 (+171 bp) | GCATCAAGTCAGCCATCAGC | CGG | GRCh38.p14 |
| IL1RN promoter 4 (+203 bp) | TGAGTCACCCTCCTGGAAAC | TGG | GRCh38.p14 |
| MYOD1 promoter 1 (+43 bp) | GCCTGGGCTCCGGGGCGTTT | AGG | GRCh38.p14 |
| MYOD1 promoter 2 (+130 bp) | GGGCCCCTGCGGCCACCCCG | GGG | GRCh38.p14 |
| MYOD1 promoter 3 (+202 bp) | CCTCCCTCCCTGCCCGGTAG | GGG | GRCh38.p14 |
| MYOD1 promoter 4 (+262 bp) | GAGGTTTGGAAAGGGCGTGC | CGG | GRCh38.p14 |
| GRM2 promoter 1 (+408bp) | GGATAGGTAAAGGGGCGCGT | GGG | GRCh38.p14 |
| GRM2 promoter 2(+326bp) | GAAGGTCACTGCGCCCCGAC | AGG | GRCh38.p14 |
| GRM2 promoter 3(+208bp) | GCGCAGAGCGAGAGCGCTCG | GGG | GRCh38.p14 |
| GRM2 promoter 4 (+250bp) | GTCTGACTATGGGGCGGAGT | GGG | GRCh38.p14 |

### Extended Data Table 3 | qPCR primer sequence[1–3]

| q-RT-PCR | | |
|---|---|---|
| gene name | Fw (5'-3') | Rv (5'-3') |
| GAPDH | GGTGGTCTCCTCTGACTTCAACA | GTTGCTGTAGCCAAATTCGTTG |
| IL1RN | GACCTTCTATCTGAGGAACAACC | CACAGGACAGGCACATCTT |
| MYOD1 | CTCCAACTGCTCCGACGGCAT | ACAGGCAGTCTAGGCTCGACAC |
| GRM2 | CGGTTCCTACAGTGATGTCTCC | TGGCTTGGAAGAAGTCAGGAGG |
| ChIP-primers | | |
| gene name | Fw (5'-3') | Rv (5'-3') |
| TSS +300bp | GTTAGAGCGTTGGGGACCTT | CACATGCAGAGAACTGAGCTG |
| TSS +200bp | TTCTCTGCATGTGACCTCCC | ACACACTCACAGAGGGTTGG |
| TSS +100bp | GCTGGGCTCCTCCTTGTACT | GCTGCTGCCCATAAAGTAGC |
| TSS -100bp | GACCTCCTGTCCTATGAGGC | TTCCTCTCTCATTTCTCAGGTG |
| TSS -200bp | AATGCTTTCAGGCTCTGGTGA | aGTGCCTGCATGGTCTCC |
| TSS -300bp | CAGGTGAACAGAGAGGTGTAAC | GGCTATTTCACCAATTTCCCTATTC |
